# Decreased Mitochondrial Bioenergetics and Function Define a Distinct Metabolic Phenotype in Healthy Sedentary Individuals Detectable Through Non-Invasive CPET

**DOI:** 10.1101/2024.08.19.608601

**Authors:** Iñigo San-Millán, Janel L. Martinez, Genevieve C. Sparagna, Angelo D’Alessandro, Davide Stefanoni, Travis Nemkov, John Hill

## Abstract

**Background:** Physical inactivity is a major contributor to cardiometabolic disease and mortality. Although mitochondrial dysfunction characterizes overt pathology, whether a distinct mitochondrial phenotype is present in apparently healthy sedentary adults remains unclear.

**Methods:** Nine sedentary (SED) and ten physically active (AC) healthy males (42 ± 14 yr) were studied. Skeletal muscle bioenergetics were assessed using high-resolution respirometry, fluxomics, metabolomics and protein expression analyses. Whole-body physiology was evaluated using cardiopulmonary exercise testing (CPET) including fat oxidation and blood lactate measurements.

**Results:** At rest, SED exhibited marked reductions in mitochondrial capacity, including Complex I (−36%), Complex II (−28%), electron transport system capacity (−34%), and ATP-synthase–coupled respiration (−30%, all p < 0.01). The most pronounced alteration was a 49% reduction in mitochondrial pyruvate carrier (MPC1) expression, which closely correlated with reduced pyruvate oxidation (−37%, p = 0.006) and lower TCA intermediates. SED also showed reduced MCT1 abundance, impaired fatty acid oxidation capacity (−32% to −35%), decreased CPT1 activity (−51%), altered cardiolipin composition and elevated ROS/O flux ratios. During exercise, SED demonstrated lower VO max (−38%), reduced fat oxidation (−35%) and higher blood lactate accumulation (>60%, p < 0.001). Mitochondrial function was strongly associated with exercise performance (r = 0.57–0.78, p < 0.01).

**Conclusions:** Healthy sedentary adults are characterized by reduced mitochondrial function characterized by decreased substrate entry and oxidation, reduced oxidative capacity and diminished metabolic flexibility. CPET-derived fat oxidation and blood lactate responses closely reflect skeletal muscle mitochondrial function, providing non-invasive physiological markers of metabolic health.

**Graphical Abstract:** Figure 1.
Schematic of skeletal muscle mitochondrion: SED side shows reduced MPC, CPT1, L4CL, and TCA flux with elevated ROS; AC side shows robust OXPHOS, fat oxidation, and lactate clearance. Arrows link to CPET as a non-invasive diagnostic tool for mitochondrial health.

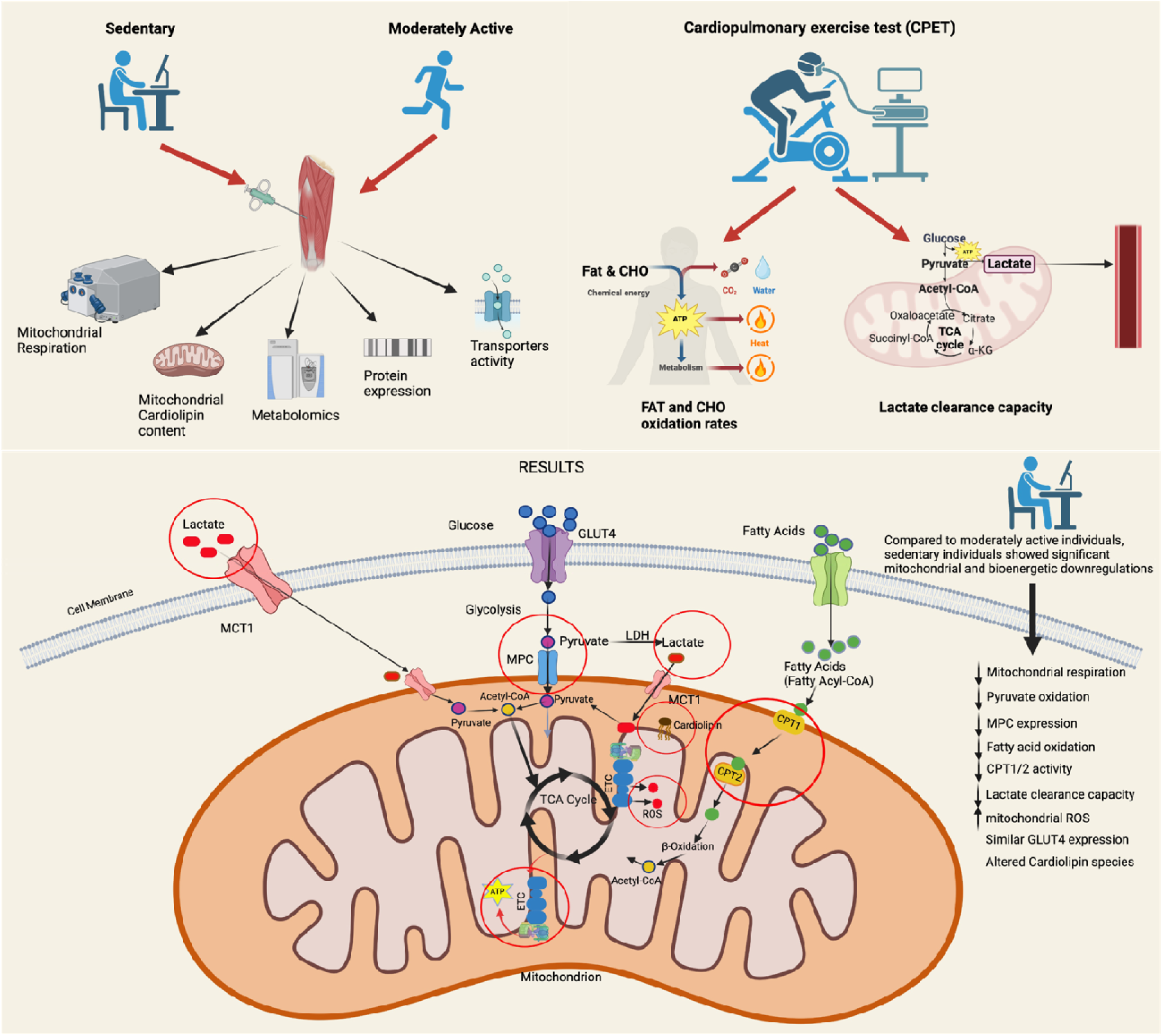

## Introduction

Physical inactivity is a global health crisis, contributing to over 5 million deaths annually through non-communicable diseases (NCDs) such as cardiovascular disease (CVD), type 2 diabetes (T2D), Alzheimer’s disease (AD), Parkinson’s disease, depression, dementia, and cancer(Ahmed et al., 2012; Berlin & Colditz, 1990; Chen et al., 2005; Colberg et al., 2016; Durstine et al., 2013; Friedenreich et al., 2021; Grill et al., 2017; Kohl 3rd, 2001; LaMonte et al., 2005; McTiernan et al., 2019; Powell et al., 1987; Rolland et al., 2008; Santos-Lozano et al., 2016; Stephen et al., 2017). Low cardiorespiratory fitness (CRF) is the leading attributable risk factor for all-cause mortality, surpassing smoking and obesity (Blair, 2009). The World Health Organization (WHO) reports that 27.5% of adults and 81% of adolescents fail to meet physical activity guidelines, projecting 500 million new NCD cases by 2030, with treatment costs of ∼US$27 billion annually and billions more in productivity losses(World Health, 2022).

Physical activity is a canonical characteristic of humans. Nevertheless, in modern societies, the normalization of the lack of physical activity has led to the perception that physical activity is an intervention even though it remains as the *modus vivendi* engrained in our genes. The reality is that becoming sedentary has been the real intervention and collateral effect of modern societies(San-Millán, 2023). This evolutionary perspective challenges the use of “healthy sedentary” individuals as controls in medical research, as they may harbor subclinical metabolic dysregulations. Skeletal muscle is responsible for ∼80-85% of glucose disposal under resting and hyperinsulinemic-euglycemic postprandial conditions (DeFronzo & Tripathy, 2009) and therefore a critical site for studying cellular metabolism, mitochondrial function and bioenergetics. Mitochondrial dysfunction is implicated in the pathogenesis of NCD’s, including T2D, CVD, AD and even cancer all characterized by impaired oxidative phosphorylation (OXPHOS) and substrate metabolism(San-Millán, 2023). Once glucose enters the cell through GLUT transporters glycolysis converts it to pyruvate which under normal flux and resting conditions should enter mitochondrial for complete oxidation through OXPHOS in the electron transport chain (ETC). The discovery of the mitochondrial pyruvate carrier (MPC) in 2012 (Bricker et al., 2012) revolutionized our understanding of glucose metabolism, highlighting pyruvate transport as a pivotal step linking glycolysis to mitochondrial ATP production. Dysregulated MPC may precede insulin resistance, offering a novel therapeutic target.

Despite the global burden of sedentarism, few studies have characterized the metabolic and bioenergetic profiles of “healthy sedentary” individuals compared to moderately active counterparts, the true evolutionary control. Early studies focused on overt disease states (e.g., T2D, obesity), but subclinical impairments in sedentary individuals could provide a preventive window. Mitochondrial respiration, substrate oxidation (carbohydrates, fatty acids, amino acids), MPC content, reactive oxygen species (ROS), and lipid composition (e.g., cardiolipin) are key indicators of cellular health. Additionally, exercise-based assessments like CPET can reveal metabolic flexibility and lactate clearance, potentially mirroring resting mitochondrial function non-invasively(San-Millan & Brooks, 2018), which offers significant possibilities for earlier diagnosis, prevention through exercise and nutrition as well as even the development of novel therapeutics.

Herein, we aimed to characterize mitochondrial and bioenergetic signatures in sedentary (SED) vs. moderately active (AC) individuals, using resting muscle biopsies (high-resolution respirometry, fluxomics, metabolomics, protein expression, cardiolipin analysis) and exercise testing (CPET, fat/carbohydrate oxidation, lactate clearance). We hypothesized that SED individuals exhibit significant mitochondrial impairments, detectable at rest and during exercise, which could be assessed non-invasively via CPET. Our findings reveal early bioenergetic decline in SED, offering insights into NCD prevention and novel therapeutic targets like MPC.

## Methods

### 2.1 Subject Recruitment

Nineteen healthy male subjects (age: 41.9 ± 13.8 years) were recruited and assigned to two groups according to their habitual physical activity levels.

- **Sedentary (SED, n=9):** No regular exercise or elevated heart rate beyond daily tasks.
- **Active (AC, n=10):** ≥150 min/week aerobic exercise for ≥6 months.

Anthropometric characteristics were as follows: the sedentary group (n = 9) exhibited a mean body weight of 90.1 ± 23.7 kg, height of 177.6 ± 5.7 cm, and body mass index (BMI) of 28.3 ± 6.0 kg·m ². The active group (n = 10) presented a mean body weight of 78.5 ± 6.0 kg, height of 181.6 ± 4.0 cm, and BMI of 23.8 ± 1.0 kg·m ².

Exclusion criteria: diagnosed diabetes (type 1 or 2), or cardiovascular risk per American College of Sports Medicine (ACSM) Risk Stratification Model. All subjects provided informed consent. The study was approved by the Colorado Multiple Institutional Review Board (17-1095).

### 2.2 Muscle Biopsies

Biopsies (∼75 mg total) from the vastus lateralis were obtained under ultrasound guidance using a Bard Monopty Disposable Core Biopsy Instrument (12 gauge × 10 cm) after local anesthesia (5 mL 1% lidocaine HCl, Pfizer) and Ethyl Chloride spray (Gebauer). Three passes yielded ∼25 mg tissue each. Samples were immediately processed fresh for respirometry or flash-frozen in liquid nitrogen and stored at -80°C. Incisions were closed with Steri-strips (3M Nexcare) and Elastoplast tape. n=10/group.

### 2.3 High-Resolution Respirometry and ROS Production

Permeabilized muscle fibers (∼10 mg) were prepared in BIOPS (10 mM Ca-EGTA, 0.1 µM free calcium, 20 mM imidazole, 20 mM taurine, 50 mM K-MES, 0.5 mM DTT, 6.56 mM MgCl, 5.77 mM ATP, 15 mM phosphocreatine, pH 7.1) with 30 µg/mL saponin for 30 min. Fibers were washed for 10 minutes at 4°C in ice-cold mitochondrial respiration medium(Gnaiger et al., 2000). Samples were blotted on filter paper, weighed, and placed in the chambers of the Oroboros O2K apparatus at 37°C containing respiration medium. Substrates were added sequentially: palmitoylcarnitine (0.2 mM) + malate (1 mM) + ADP (4 mM) for long-chain fatty acid oxidation; octanoylcarnitine (0.4 mM) for medium-chain; pyruvate (5 mM) for complex I via pyruvate dehydrogenase; glutamate (10 mM) for complex I independent of PDH; succinate (10 mM) for complex II; FCCP (1 µM) for maximal ETS capacity; rotenone (2 µM) for complex II; antimycin A (5 µM) for residual oxygen consumption (ROX). ROS was measured via Amplex Red (5 units/mL SOD, 1 unit/mL horseradish peroxidase, 0.1 µM/step H O calibration). O2 flux normalized to tissue wet weight. n=10/group.

**Figure 2.**
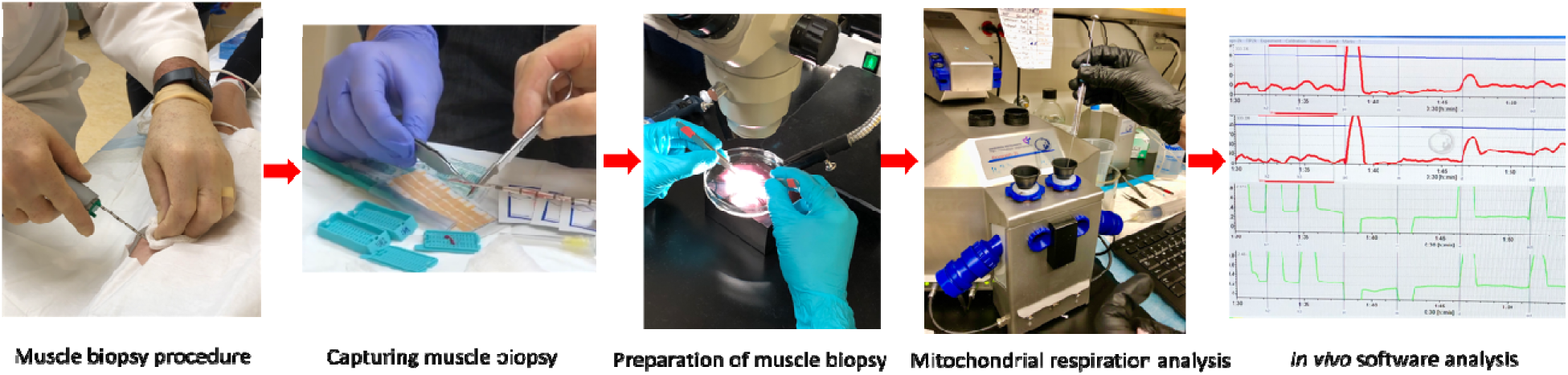
Muscle biopsy procedure included extraction, mitochondrial homogenization, and mitochondrial respiration analysis through Oroboros O2K.

### 2.4 Protein Expression

Skeletal muscle homogenates were analyzed via capillary-based Western (Jess, ProteinSimple). Antibodies: GLUT4 (Abcam ab654, 1:1000), LDHA (Cell Signaling #2012, 1:500), LDHB (Cell Signaling #8176, 1:500), MPC1 (Abcam ab176559, 1:800). Samples were denatured (95°C, 5 min) with fluorescent master mix (1:4). Protein peak areas were normalized to total protein using the Compass for Simple Western software (ProteinSimple, CA). Antibody specificity was validated via molecular weight controls. n=9-10/group (minor sample loss in 1 SED).

### 2.5 Carnitine Palmitoyltransferase I and II Activity

CPT1 and CPT2 activity was quantified in muscle homogenates using a ¹ C-carnitine radioassay (20). CPT1: plasma membrane permeabilized, measuring palmitoylcarnitine production from palmitoyl-CoA. CPT2: mitochondrial inner membrane permeabilized with malonyl-CoA to inhibit CPT1. n=9-10/group.

### 2.6 Skeletal Muscle Isotope Tracing

Fresh tissue (∼10 mg) was incubated (37°C, 30 min) in Krebs-Ringer bicarbonate solution with 2 mM ¹³C-pyruvate (CLM-2440-PK, Cambridge Isotope Laboratories) or 2 mM ¹³C-lactate (CLM-1577-PK). Samples were centrifuged (2000g, 10 min), flash-frozen, and stored at -80°C. n=9-10/group.

### 2.7 Mass Spectrometry-Based Metabolomics

Muscle tissue was extracted in methanol:acetonitrile:water (5:3:2) at 30 mg/mL, vortexed (4°C, 30 min), and centrifuged (10,000g, 10 min, 4°C). Supernatants were analyzed via UHPLC-MS (Vanquish/Q Exactive, Thermo Fisher) over a Kinetex C18 column (2.1 × 150 mm, 1.7 µm, Phenomenex). Mobile phases: positive ion (water + 0.1% formic acid; acetonitrile + 0.1% formic acid); negative ion (water:acetonitrile 95:5 + 1 mM ammonium acetate; acetonitrile:water 95:5 + 1 mM ammonium acetate). Gradient: 5% to 95% B over 1 min, hold at 95% B for 2 min (400 µL/min, 45°C). Full MS mode (60-900 m/z, 70,000 resolution). Data converted to mzXML (Mass Matrix), processed in El-MAVEN for metabolite assignments and isotopologue distributions, corrected for natural isotope abundances. Median-normalized, range-scaled data analyzed via MetaboAnalyst 6.0 (PLS-DA, hierarchical clustering). n=10/group (292 metabolites quantified).

**Figure 3.**
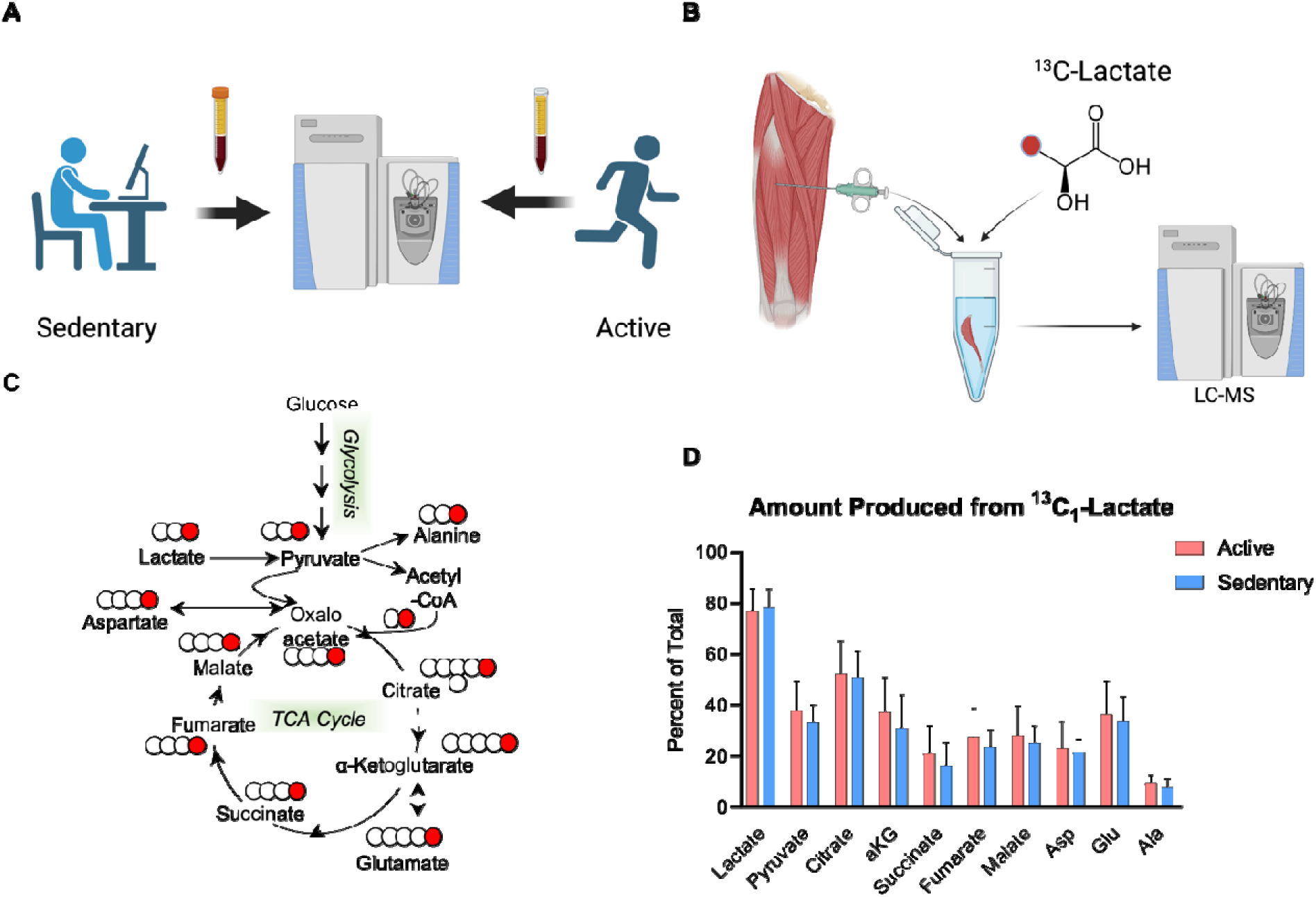
Ex Vivo Skeletal Muscle Isotope Tracing Analysis. (A) Skeletal muscle was isolated from 10 sedentary and 13 active individuals. (B) Isolated tissue was incubated with ^13^C_1_-lactate for 30 minutes in a physiological buffer at 37C. (C) Isotope tracing was targeted to metabolites in the TCA Cycle and related transaminase products. (D) Isotope enrichment is reported as a percentage of the total pool in each sample.

### 2.8 Cardiolipin Quantification

Lipid extracts from homogenized muscle (PBS) were analyzed via LC/MS (API 4000, Sciex) using normal phase solvents(Sparagna et al., 2005). Total cardiolipin (CL) was the sum of seven dominant species (m/z 1422, 1446, 1448, 1450, 1470, 1472, 1474). L4CL percentage calculated. n=9-10/group.

### 2.9 Graded Exercise Assessment

Subjects consumed >50% kcal as carbohydrates the night before/day of testing (verified via food logs). Exercise was performed on a cycle ergometer (KICKR Smart Trainer, Wahoo, USA). Warm-up: <65W for 10 min. Protocol: start at 75W, increase by 25W every 10 min until volitional exhaustion (inability to maintain 60 rpm). All subjects reached ≥150W. n=9-10/group.

### 2.10 Gas Exchange Measurements

VO, VCO, and respiratory exchange ratio (RER = VCO /VO) were measured via ParvoMedics TrueOne 2400 (Sandy, UT, USA), averaged over 15-s intervals. n=9-10/group.

### 2.11 Fat and Carbohydrate Oxidation Rates

Calculated per Frayn(Frayn, 1983):

- FATox (g·min ¹) = 1.67 VO (L·min ¹) – 1.67 VCO (L·min ¹)
- CHOox (g·min ¹) =4.55 VCO (L·min ¹) – 3.21 VO (L·min ¹)

n=10/group.

### 2.12 Lactate Concentration Measurement

Capillary blood from the earlobe was analyzed for L-lactate (Lactate Plus Meter, Nova Biomedical, USA) at each stage’s end. Heart rate (Polar S725x) and perceived exertion were monitored. n=9-10/group.

### 2.13 Statistical Analysis

Data analyzed in GraphPad Prism 9.2.1. Independent t-tests with Shapiro-Wilk normality test; Cohen’s d for effect sizes. Bonferroni correction for multiple comparisons (α=0.05/num tests). Pearson correlations for rest-exercise relationships; Spearman’s used if non-normal. Data: mean ± SD; p<0.05 significant, p<0.1 as trending. Power analysis (80% power, α=0.05) estimated n=9-10/group sufficient for large effects (d>1.0).

## Results

### 3.1 Mitochondrial Respiration and Oxidative Capacity

At rest, sedentary individuals (SED) exhibited marked deficits across multiple components of the mitochondrial electron transport system (ETS) when compared with active individuals (AC) (Fig. 4A–D). Oxygen flux through Complex I was reduced by 36% (p = 0.008, d = 1.8), and Complex II–linked respiration was 28% lower (p = 0.042, d = 1.2), indicating attenuated substrate entry and electron transfer capacity through both NADH- and FADH-dependent pathways. The deficit extended to total ETS capacity which was 34% lower in SED (p = 0.007, d = 1.7) and to ETS coupled to ATP synthase (P_ATP) which was reduced by 30% (p = 0.009, d = 1.5). Together, these data demonstrate a generalized impairment of oxidative phosphorylation efficiency and maximal mitochondrial respiratory capacity in sedentary skeletal muscle.

**Figure 4.**
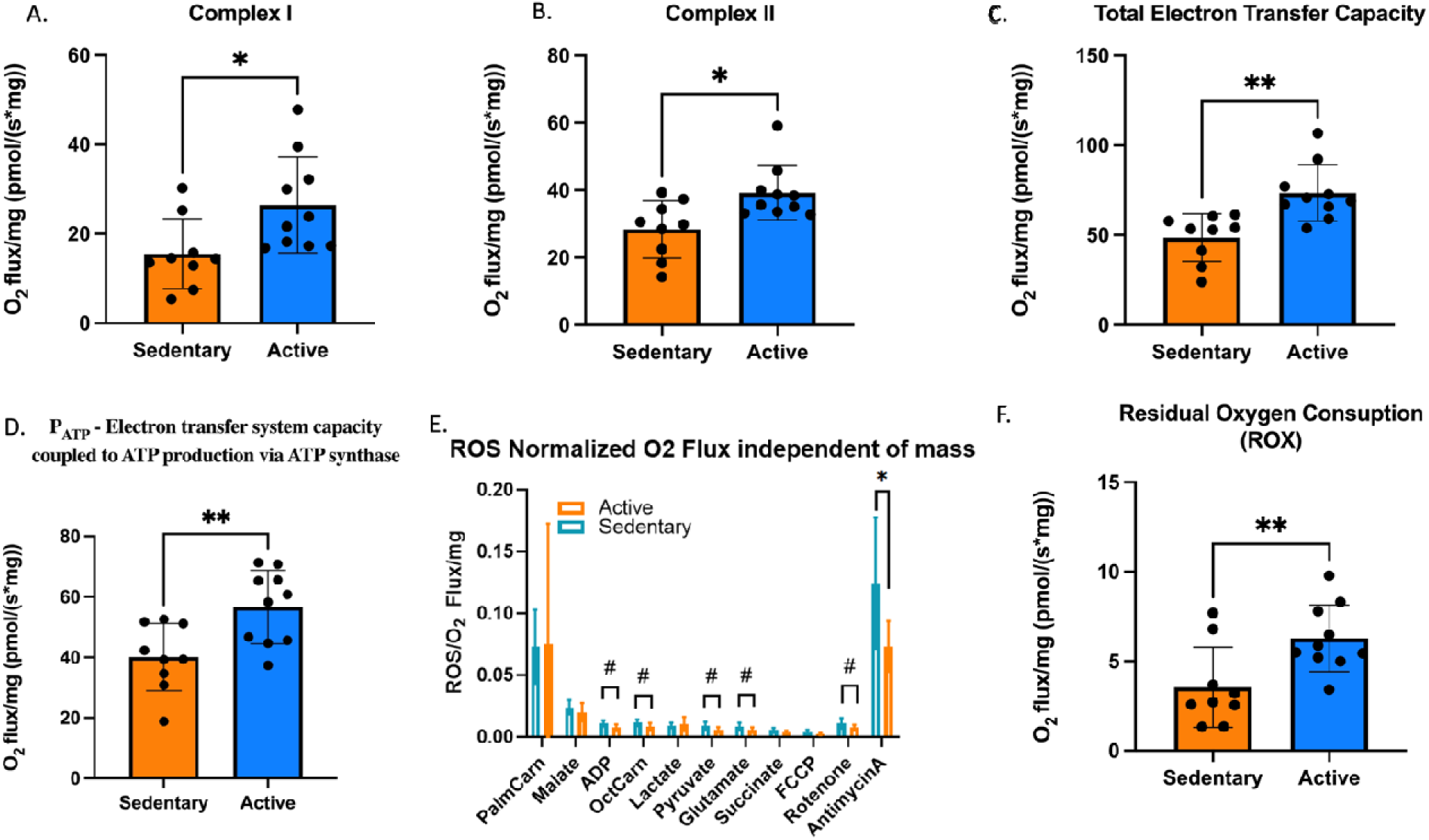
Differences in A) mitochondrial complex I capacity. B) complex II capacity. C) total electron transfer system capacity (maximal respiration). D) electron transfer system capacity coupled to ATP production via ATP synthase between sedentary and active individuals. E) Differences in reactive oxygen species (ROS) production normalized to O_2_ flux independent of mass in sedentary and active individuals. F) Differences in residual oxygen consumption (ROX) due to oxidative side reactions that continue after the inhibition of the ET-pathway. *p<0.05, **p<0.01.

### 3.2 Reactive Oxygen Species Production

Reactive oxygen species (ROS) production, when normalized to O flux, revealed further distinctions between groups (Fig. 4E–F). While absolute ROS emission tended to be higher in AC, normalization to O flux indicated greater oxidative stress burden in SED, particularly following antimycin A inhibition (p = 0.048, d = 1.1), with similar upward trends observed during ADP-, octanoylcarnitine-, pyruvate- and glutamate-driven respiration stages (p < 0.10). These findings point to a lower redox efficiency in sedentary muscle, where a greater fraction of electrons is diverted toward ROS generation rather than productive respiration.

By contrast, the higher absolute ROS levels in active individuals likely reflect physiological eustress through a controlled redox signaling milieu accompanying elevated metabolic flux and mitochondrial turnover. Thus, despite greater respiratory activity, the active phenotype maintains effective coupling between electron transport and oxidative metabolism, whereas sedentary muscle displays reduced respiratory control and elevated ROS per unit O consumed (Fig. 4-E).

Furthermore, residual oxygen consumption (ROX), the non-phosphorylating respiration that persists after inhibition with antimycin A, was significantly higher in active individuals (p = 0.007, d = 1.4) (Fig. 4-F). ROX reflects oxygen use not linked to ATP synthesis, associated with baseline mitochondrial maintenance and proton leak. The higher ROX observed in active muscle suggests greater mitochondrial density and metabolic turnover, indicative of a more dynamic and resilient oxidative

### 3.3 Protein Expression and Enzyme Activity

Skeletal muscle expression of GLUT4, LDHA, and LDHB was similar between active (AC) and sedentary (SED) individuals (Fig. 5A, C–D), indicating comparable basal capacity for glucose uptake and cytosolic lactate interconversion. Despite similar GLUT4 content, MPC1 expression (the mitochondrial pyruvate carrier subunit critical for pyruvate transport into the matrix) was markedly lower in SED, reduced by 49% relative to AC (p = 0.005, d = 2.1). This deficit strongly correlated with diminished pyruvate-driven respiration, suggesting a functional limitation in the entry of glycolytic carbon into oxidative pathways.

**Figure 5.**
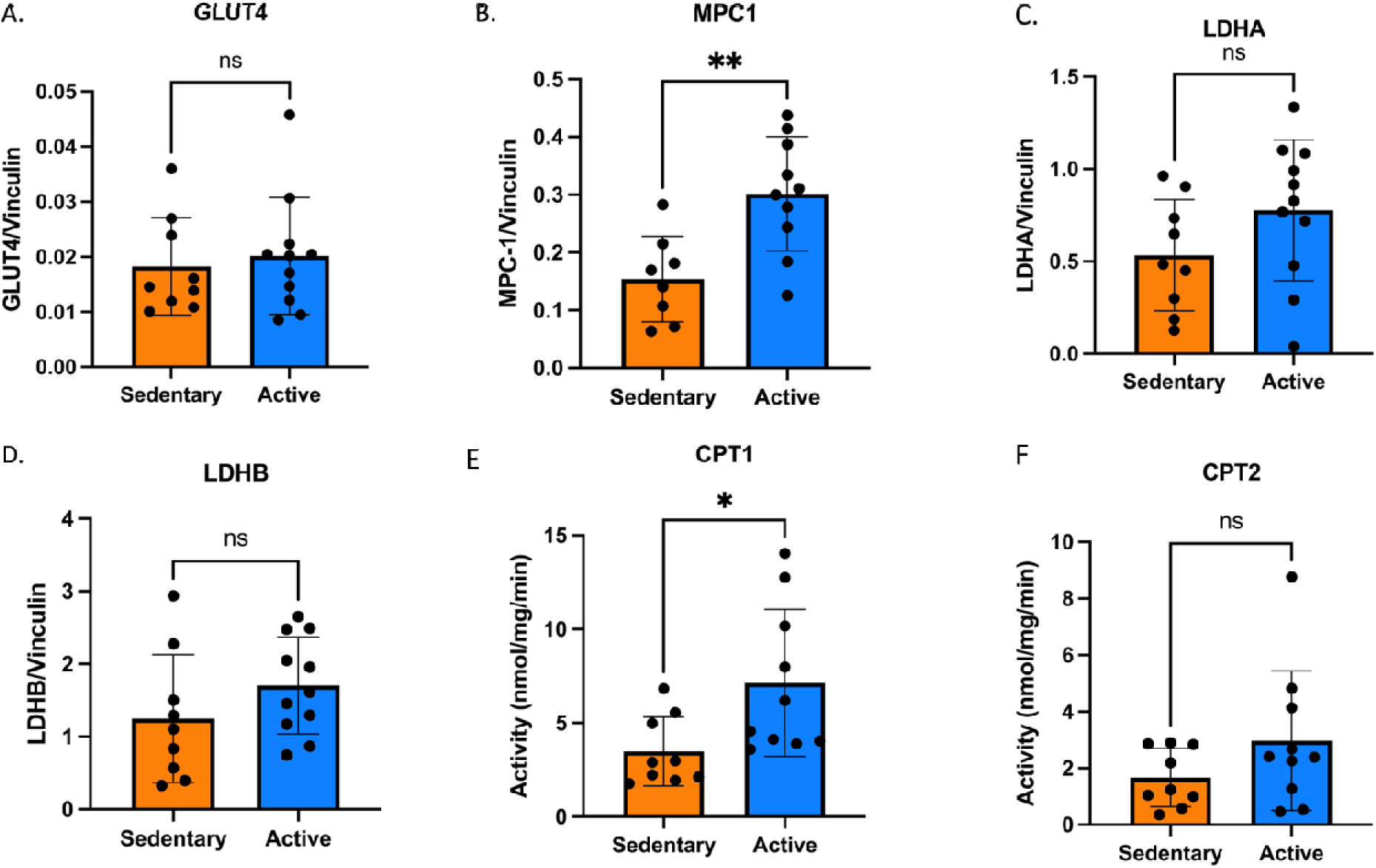
Differences in protein expression between SED and ACT individuals of A) GLUT4, B) MPC1, C) LDH_A_, D) LDH_B_, E) CPT1 and F) CPT2 between sedentary and active individuals. E). *p<0.5, ** p<0.01.

In parallel, CPT1 activity, the rate-limiting enzyme for long-chain fatty acid transport across the outer mitochondrial membrane, was 51% lower in SED (p = 0.038, d = 1.4), whereas CPT2 activity showed a downward trend without reaching significance (p = 0.16) (Fig. 5E–F). Together, these findings indicate that sedentary muscle exhibits convergent impairments in both pyruvate and fatty acid transport systems, compromising the ability to flexibly switch between carbohydrate and lipid substrates for mitochondrial oxidation and therefore, affecting metabolic flexibility.

### 3.4 Cardiolipin Composition

Mitochondrial phospholipid analysis revealed clear group differences in cardiolipin content and remodeling profiles (Fig. 6A–E). Total cardiolipin and its dominant molecular species, tetralinoleoyl-cardiolipin (L4CL), were significantly lower in sedentary individuals (p = 0.007 and p = 0.006, respectively; d ≈ 1.5–1.6). In contrast, the proportion of monolysocardiolipin (MLCL), a remodeling intermediate, was modestly higher in active individuals (p = 0.049), resulting in a slightly elevated MLCL:CL ratio. These compositional differences suggest that active muscle maintains greater cardiolipin abundance and enrichment in the mature L4CL species, which is essential for optimal organization of respiratory supercomplexes and efficient electron transfer. Conversely, the lower total and tetralinoleoyl cardiolipin content observed in sedentary muscle implies reduced inner-membrane structural integrity and less efficient OXPHOS organization, consistent with the observed declines in ETS capacity and ATP-linked respiration.

**Figure 6.**
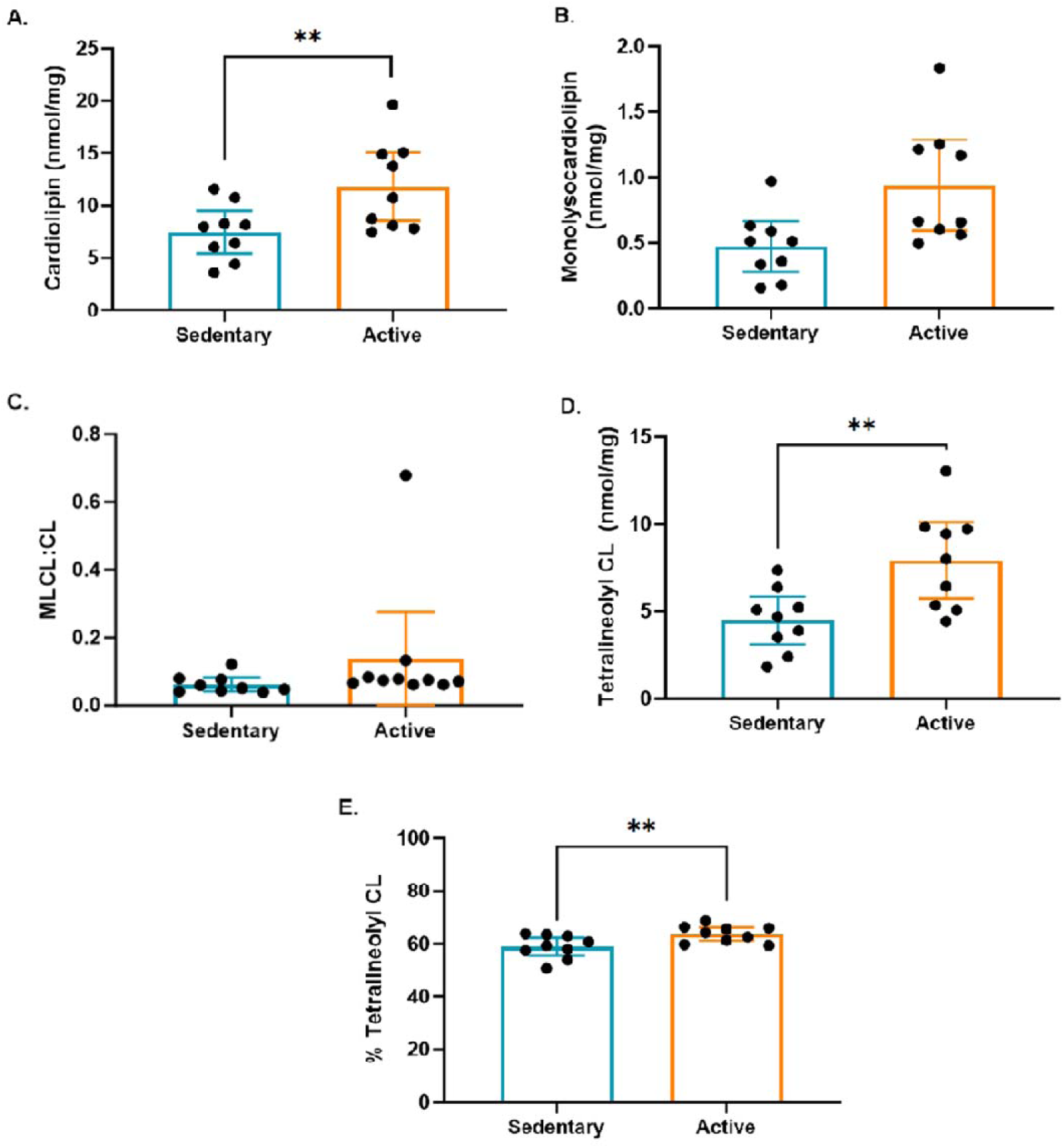
Cardiolipin content and molecular composition in skeletal muscle. (A) Total cardiolipin, (B) monolysocardiolipin, (C) MLCL:CL ratio, (D) tetralinoleoyl cardiolipin (L4CL), and (E) percent L4CL. Both total cardiolipin and L4CL were significantly higher in active individuals (p < 0.01), whereas MLCL showed a modest increase (*p < 0.05). Bars represent mean ± SD. *p < 0.05, \**p < 0.01; ns = not significant*.

### 3.5 Substrate-Specific Mitochondrial Oxidation

Substrate-dependent respiration revealed a consistent deficit across multiple oxidative pathways in sedentary individuals (Fig. 7A–D). Pyruvate-supported O flux was 37% lower in SED compared with active individuals (p=0.006, d=1.9), indicating reduced capacity for mitochondrial oxidation of glycolytic carbon. This limitation aligns with the markedly lower MPC1 expression observed in SED muscle suggesting a bottleneck at the level of mitochondrial pyruvate transport and thus, a critical gateway for coupling cytosolic glycolysis to mitochondrial ATP production.

**Figure 7.**
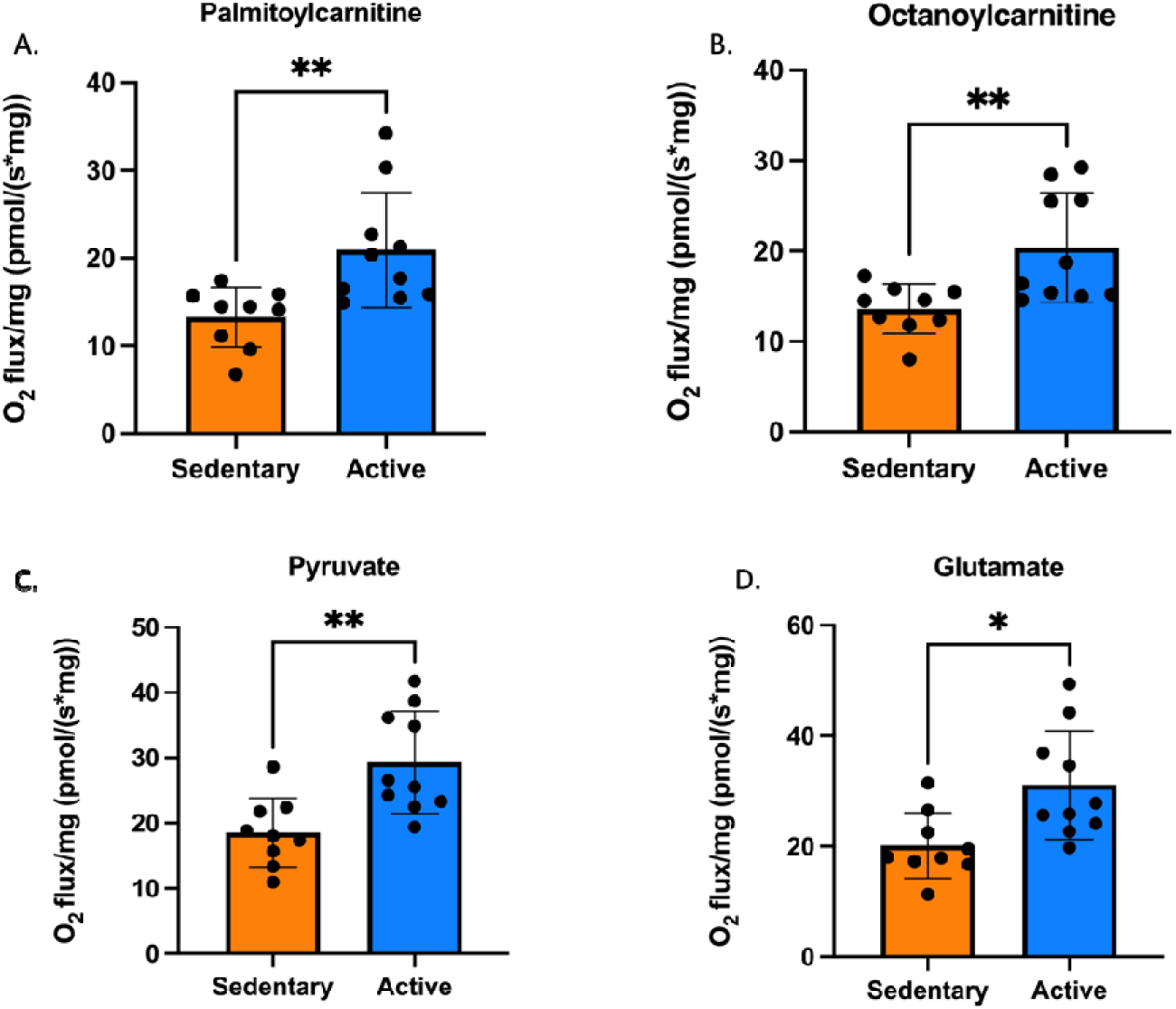
Differences in O_2_ flux between sedentary and active individuals following saturating additions of A) palmitoylcarnitine, indicating long-chain fatty acid oxidation, B) octanoylcarnitine, indicating medium-chain fatty acid oxidation C) pyruvate, indicating carbohydrate-derived oxidation through complex I following transport into mitochondria via mitochondrial pyruvate carrier (MPC) and D) glutamate through complex I independent of pyruvate dehydrogenase complex). *p<0.5, ** p<0.01.

Similarly, long-chain fatty acid oxidation using palmitoylcarnitine and medium-chain oxidation with octanoylcarnitine were lower in SED by 35% and 32%, respectively (both p = 0.008, d ≥ 1.5). These findings point to a parallel deficit in lipid-derived electron flux, consistent with the reduced CPT1 activity reported earlier. Glutamate-driven Complex I respiration was also reduced by 36% (p=0.008, d=1.5), reinforcing the notion of a global decline in oxidative substrate utilization.

Collectively, these results reveal that sedentary skeletal muscle exhibits a multifaceted impairment in substrate oxidation, encompassing both carbohydrate- and lipid-derived pathways. The concurrent reductions in MPC1 expression and pyruvate oxidation capacity underscore a key mechanistic link between limited mitochondrial substrate transport and the diminished metabolic flexibility characteristic of the sedentary phenotype.

### 3.6 Metabolomic Profiling and [¹³CD]-Lactate Tracing

Multivariate analysis of the untargeted metabolomic data using PLS-DA (Fig. 8A) demonstrated clear discrimination between sedentary (SED) and active (AC) individuals, with distinct group clustering and minimal overlap, indicating a robust separation based on metabolic phenotype. The volcano plot (Fig. 8B) and heatmap (Fig. 8C) further highlighted differential abundance patterns between groups.

**Figure 8.**
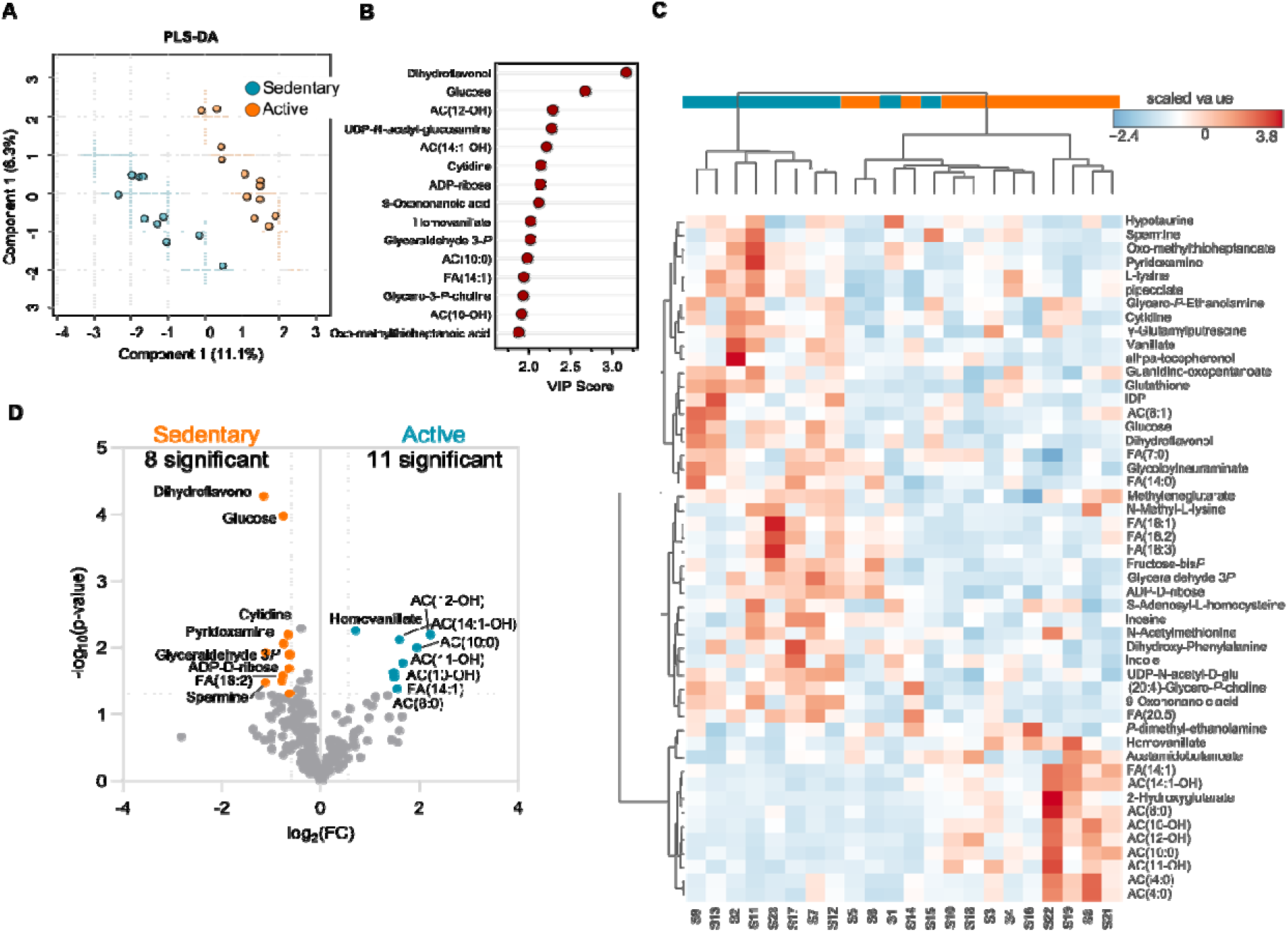
Untargeted metabolomics and [¹³C]-lactate tracing in skeletal muscle. (A) PLS-DA plot showing distinct separation between sedentary and active individuals.(B) Volcano plot and (C) heatmap illustrating differential metabolite abundance, with higher glycolytic intermediates in sedentary muscle and elevate acylcarnitines and TCA metabolites in active muscle. (D) [¹³C]-lactate tracing demonstrates reduced labeling of citrate and malate in sedentary individuals (–40% and –35%, p<0.01), indicating reduced pyruvate flux into the TCA cycle and impaired mitochondrial integration of lactate-derived carbon.

SED muscle displayed central metabolic accumulation of glycolytic and upstream carbohydrate intermediates, including glucose-6-phosphate, fructose-1,6-bisphosphate, 3-phosphoglycerate, and phosphoenolpyruvate, along with elevated lactate and pyruvate, consistent with enhanced cytosolic glycolytic flux and reduced oxidative disposal. In contrast, AC muscle exhibited higher levels of short- and medium-chain acylcarnitines (C6–C12 species), succinate and fumarate, reflecting greater lipid mobilization and mitochondrial TCA activity.

Stable-isotope tracing with [¹³C]-lactate (Fig. 8D) revealed markedly lower labeling of citrate (–40%) and malate (–35%) in SED muscle (both p < 0.01), indicating reduced pyruvate entry into the TCA cycle. This attenuation closely parallels the MPC1 deficit and diminished pyruvate-supported respiration described earlier, confirming a functional limitation in lactate-derived carbon oxidation.

Overall, these findings define a metabolic signature of inactivity characterized by glycolytic metabolite accumulation and constrained pyruvate–TCA coupling, contrasting with the broader substrate oxidation and acylcarnitine enrichment observed in active skeletal muscle.

### 3.7 Exercise Performance

The exercise results revealed further significant differences between AC and ED. AC displayed higher aerobic fitness and oxidation flexibility. Absolute and relative VO2max were 31% and 38% greater respectively by (both p < 0.0001, d > 2.2) (Fig-9A-B). Maximal and relative absolute power output were 35% and 42% higher respectively (both p < 0.0001) (Fig 9-C-D). Blood lactate during moderate stages at 125W and 150W respectively, was higher in SED (p < 0.001, d ≈ 1.5) (Fig-9E-F).

**Figure 9.**
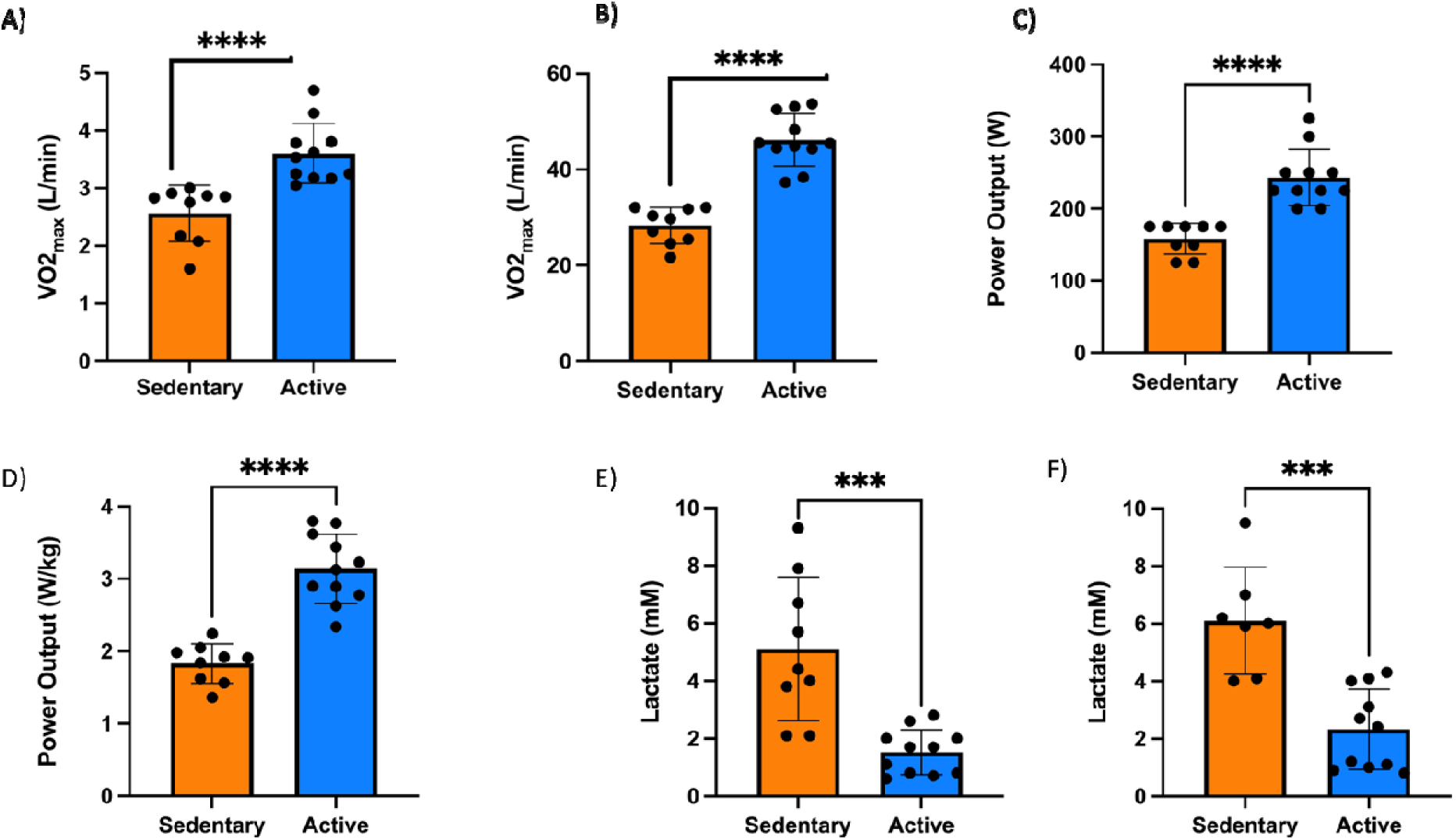
VO2max and power output and blood lactate differences between both AC and SED groups. ***p<0.001, **** p<0.0001.

### 3.8 Substrate Oxidation and Lactate Responses During Incremental Exercise

Figure 10 (Panels A–F) illustrates the distinct metabolic responses between active and sedentary individuals across increasing power outputs. In active individuals (Panels A, C, E), fat oxidation (FATox) progressively rises at low-to-moderate workloads and peaks before declining at higher intensities as carbohydrate oxidation (CHOox) and blood lactate concentrations increase. FATox reaches ∼0.35–0.4 g·min ¹ at ∼150 W, with lactate remaining below 1 mM throughout this range (r = –0.99, p < 0.001), reflecting a high capacity for mitochondrial substrate oxidation and efficient redox balance. As workload increases beyond this point, CHO oxidation rises sharply (r = 0.99, p < 0.001), paralleling the gradual elevation in lactate levels. This pattern demonstrates a well-preserved metabolic flexibility, characterized by the ability to transition efficiently from predominant lipid oxidation to carbohydrate utilization as exercise intensity increases.

**Figure 10.**
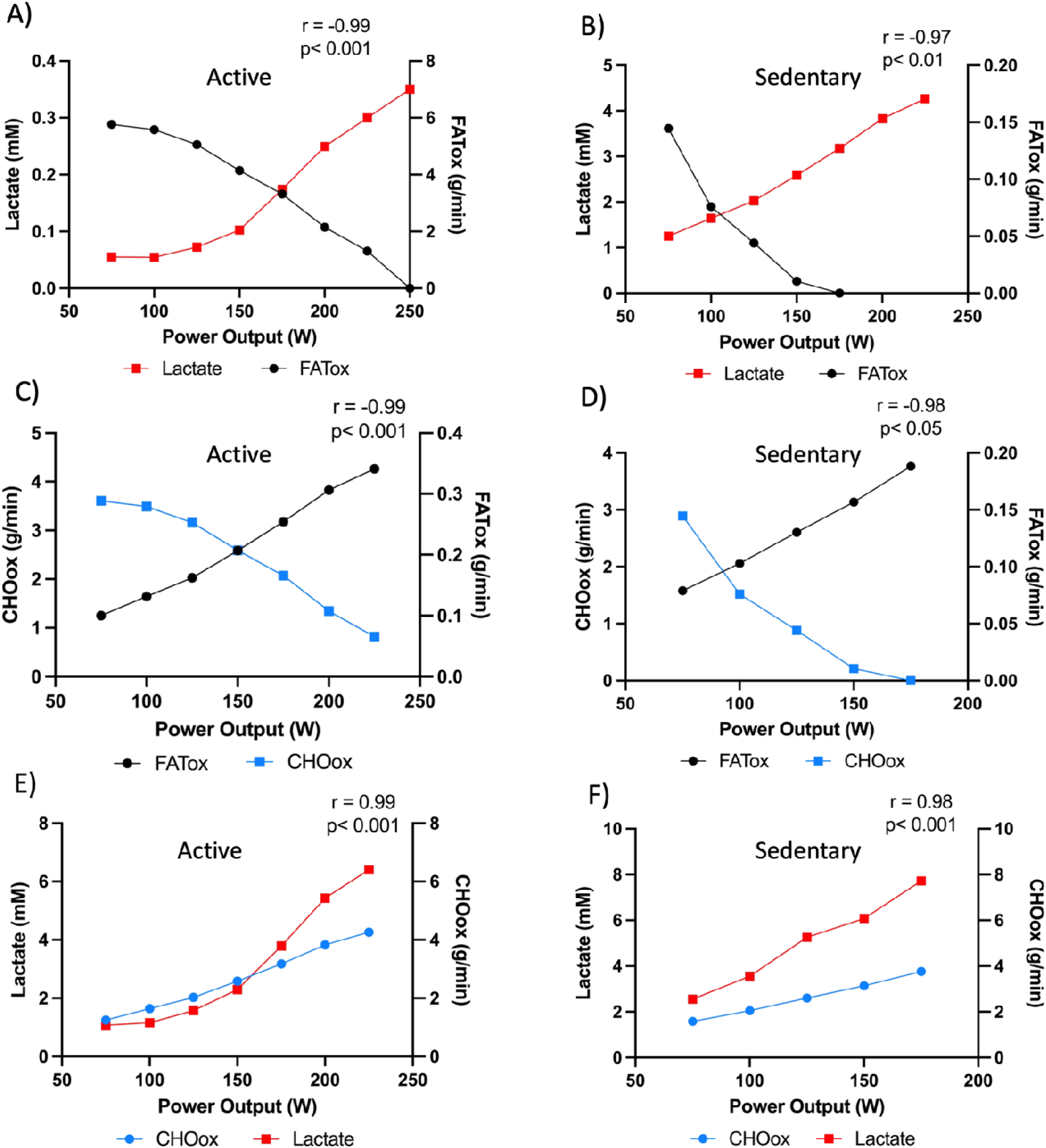
Substrate oxidation and lactate dynamics during incremental exercise in active and sedentar individuals. (A, C, E) Active individuals show progressive increases in fat oxidation (FATox) at low intensities followed by a rise in carbohydrate oxidation (CHOOx) and lactate at higher workloads. (B, D, F) Sedentary individuals display lower FATox and earlier carbohydrate dependence, accompanied by a steep lactate rise even at moderate workloads. Correlation coefficients (r) reflect strong reciprocal relationships between FATox and lactate (A–B) and between CHOOx and FATox (C–D). These relationships illustrate preserved metabolic flexibility in active subjects and impaired substrate switching in sedentary counterparts, with lactate serving as a surrogate of mitochondrial oxidative efficiency.

Conversely, in sedentary individuals (Panels B, D, F) the metabolic profile is markedly altered. FATox remains substantially lower across all intensities, peaking near 0.15–0.2 g·min ¹ and declining rapidly with rising power output while lactate accumulates steeply from the earliest workloads (r = –0.97, p < 0.01). CHO oxidation dominates even at low intensities, increasing linearly with power output, indicative of an early metabolic inflexibility. The elevated lactate concentrations at submaximal workloads suggest reduced mitochondrial pyruvate oxidation capacity and earlier increased glycolytic flux consistent with impaired oxidative metabolism.

The strong reciprocal correlations between lactate and FATox (r = –0.97 to –0.99) across groups emphasize the tight coupling between lactate production and the suppression of fat oxidation. These findings position lactate not merely as a by-product of glycolysis but as a functional biomarker of substrate shift and mitochondrial efficiency. In metabolically flexible, active individuals, efficient pyruvate transport and oxidation within the mitochondria limit lactate accumulation and maintain high lipid utilization. In contrast, sedentary individuals exhibit an earlier glycolytic reliance and lactate accumulation that reflect a bottleneck at the mitochondrial pyruvate carrier and reduced oxidative phosphorylation capacity.

Together, these results visualize metabolic flexibility as the dynamic intersection of fat and carbohydrate oxidation, with lactate serving as an integrated systemic marker of this shift. Active individuals maintain oxidative dominance over a wide intensity range, whereas sedentary individuals exhibit a constrained oxidative phenotype and early transition to glycolytic metabolism.

### 3.9 Rest–Exercise Correlations and Diagnostic Translation

Across participants, strong associations appeared between resting mitochondrial respiration and exercise-derived substrate oxidation. Resting fatty acid–supported oxygen flux, measured using palmitoylcarnitine and octanoylcarnitine, correlated robustly with in vivo fat oxidation rates during exercise (r = 0.71, p < 0.001), as shown in Figure 11. Similarly, total electron transfer system (ETS) capacity and ATP production at rest correlated with whole-body FATox during exercise (r = 0.63–0.70, p < 0.01), underscoring the translational link between cellular respiratory competence and systemic metabolic flexibility.

**Figure 11.**
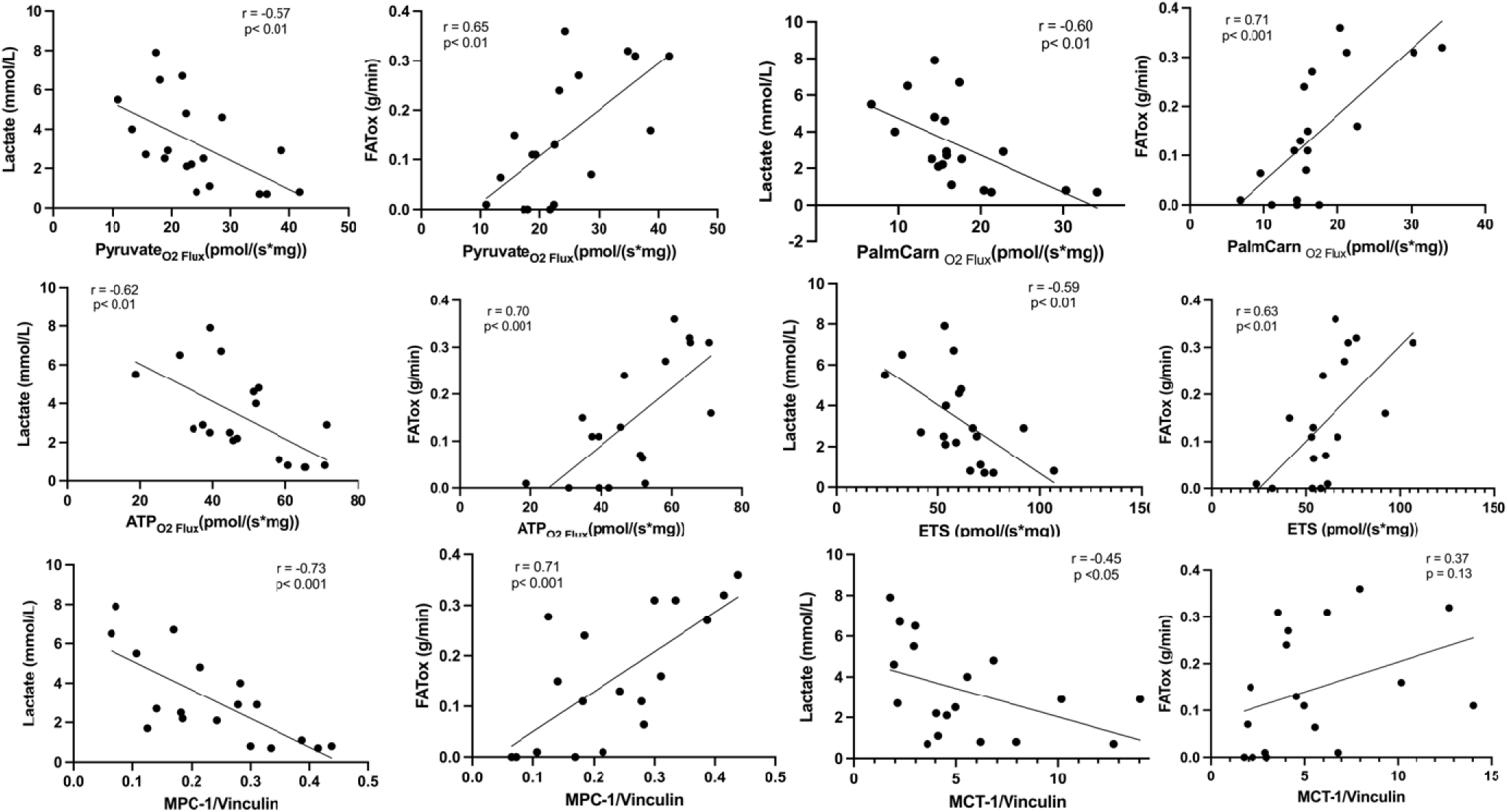
Correlations between resting mitochondrial respiration and exercise metabolic parameters. Scatterplots illustrate relationships between substrate-specific O fluxes (pyruvate, palmitoylcarnitine, octanoylcarnitine), electron transfer system capacity, and in vivo lactate and FATox responses.

Resting pyruvate-supported oxygen flux provided an additional layer of insight. Pyruvate oxidation correlated with in vivo FATox during exercise (r = 0.65, p < 0.01). Furthermore, MPC correlated with FATox during exercise (r = 0.71, p < 0.001) highlighting the central role of mitochondrial pyruvate handling in determining the transition between fat and carbohydrate utilization in vivo. Participants with higher pyruvate-supported flux at rest, reflecting greater MPC-dependent pyruvate transport, consistently showed lower lactate accumulation during exercise and a delayed shift toward carbohydrate reliance.

Maximal oxygen uptake (VO max) also correlated with ETS (r = 0.57, p < 0.01) and with combined ETS plus ATP synthase–linked capacity (r = 0.59, p < 0.01), indicating that higher electron transport and coupling efficiency at the muscle level manifest as superior aerobic performance.

Inverse correlations were observed between blood lactate concentrations during exercise and resting pyruvate-supported O flux (r = –0.57, p < 0.01), palmitoylcarnitine oxidation (r = – 0.60, p < 0.01), and MPC1 transporter (r = –0.73, p < 0.001),. Lower mitochondrial oxidative capacity at rest, particularly reduced MPC-linked pyruvate flux, was associated with earlier lactate accumulation in vivo. This pattern confirms that impaired mitochondrial pyruvate oxidation shifts reliance toward glycolysis, elevates lactate production, and narrows the metabolic flexibility window.

Together, these data provide an integrative framework linking cellular bioenergetics with systemic exercise physiology. They support the use of CPET-derived measures such as FATox, and carbohydrate oxidation rates, crossover points and lactate dynamics as non-invasive diagnostic tools to infer mitochondrial health and metabolic flexibility.

### 3.10 ¹³C-Lactate Flux and Mitochondrial Coupling

During the initial phase of sample collection, a small number of tissue specimens were lost or yielded insufficient material for all assays due to handling limitations, resulting in slightly reduced sample size for certain correlations. Nevertheless, the relationships observed were strong and internally consistent, supporting their robustness.

Lactate flux, assessed through ¹³C-lactate tracing, showed tight coupling with multiple indices of mitochondrial oxidative function (Fig. 12). The flux of labeled lactate into mitochondrial respiration correlated positively with pyruvate-supported O flux (r = 0.89, p < 0.001, *d* = 2.3) and MPC1 expression (r = 0.73, p < 0.05, *d* = 1.4), reinforcing the dependence of lactate oxidation on efficient pyruvate transport into the mitochondrial matrix. Similarly, lactate flux correlated with palmitoylcarnitine-supported respiration (r = 0.95, p < 0.0001, *d* = 3.1), octanoylcarnitine oxidation (r = 0.79, p < 0.0001, *d* = 1.7), ATP synthase–linked O flux (r = 0.81, p < 0.01, *d* = 1.8), and total ETS capacity (r = 0.84, p < 0.01, *d* = 1.9). All associations demonstrated large effect sizes (*d* > 1.2), underscoring both their biological and statistical relevance and confirming that lactate oxidation is a fully integrated component of mitochondrial respiratory function.

**Figure 12.**
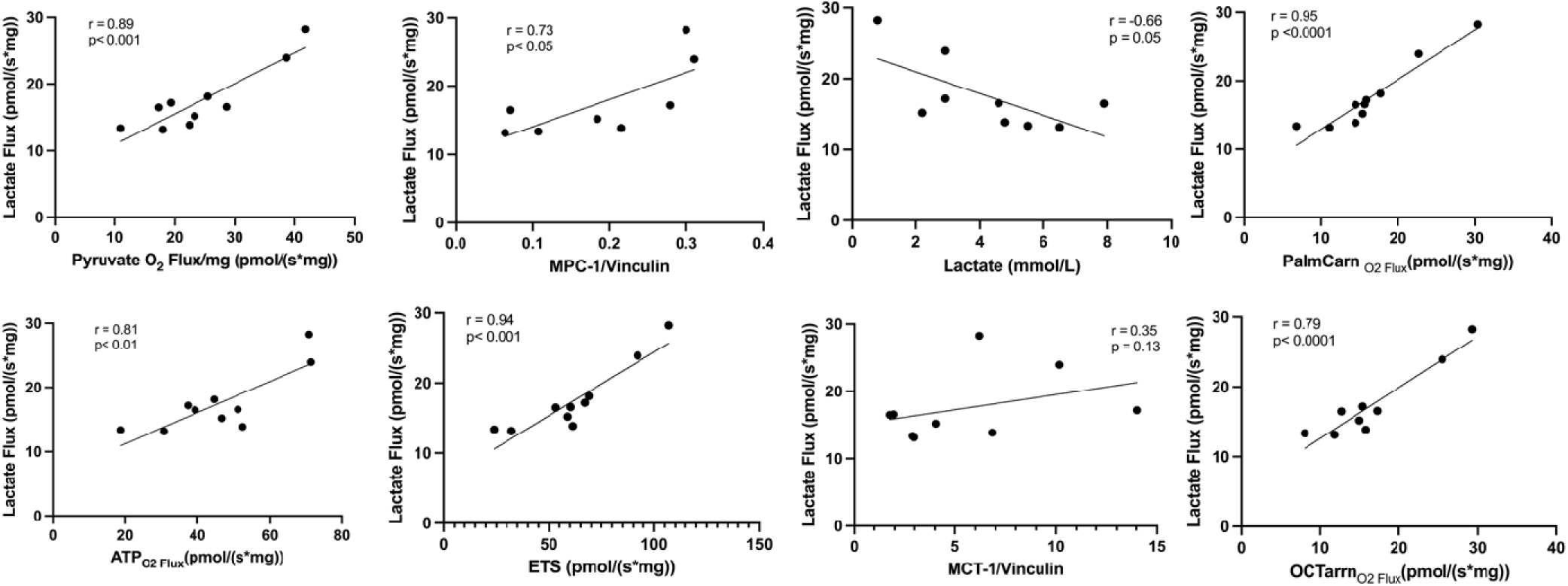
Correlations between ¹³C-lactate flux and mitochondrial functional parameters. Scatterplots show associations between lactate-derived oxidative flux and pyruvate-supported O flux, MPC1 expression, fatty-acid oxidation (palmitoylcarnitine and octanoylcarnitine), ATP-synthase–linked O flux, and total electron-transfer-system (ETS) capacity. Strong positive correlations were observed across all parameters (r = 0.73–0.95, p < 0.01, Cohen’s *d* = 1.4–3.1), indicating close coupling between lactate utilization and mitochondrial oxidative capacity. An inverse relationship was found between blood lactate concentration and lactate flux (r = – 0.66, p = 0.05), whereas total MCT1 expression was not significantly related (r = 0.35, p = 0.13). Data suggest that mitochondrial, rather than sarcolemmal, MCT1 activity governs lactate oxidation within the mitochondria. Bars and regression lines represent mean ± SD; *p* < 0.05, p < 0.01.

An inverse association was observed between blood lactate concentration and lactate flux (r = – 0.66, p = 0.05, *d* = 1.2), consistent with enhanced mitochondrial clearance capacity in individuals with higher oxidative function. No significant relationship was detected between total MCT1 expression and lactate flux (r = 0.35, p = 0.13, *d* = 0.6). This suggests that the regulation of lactate oxidation is governed primarily by mitochondrial MCT1 (not measured in the study), which facilitates intramitochondrial lactate transport and coupling to the MPC1–LDH complex, rather than by sarcolemmal MCT1 abundance measured in whole-muscle homogenates.

Taken together, these findings indicate that higher mitochondrial lactate flux, representing the dynamic utilization of lactate-derived carbon, is strongly aligned with the key determinants of oxidative metabolism, including pyruvate transport, fatty acid oxidation and ETS coupling. The magnitude of these associations supports the interpretation of lactate as both a metabolic substrate and a quantitative marker of mitochondrial efficiency.

## Discussion

This study reveals that “healthy” sedentary individuals harbor profound mitochondrial and bioenergetic impairments compared to moderately active counterparts, redefining the assumption that sedentary equals healthy. Reduced ETS capacity, MPC1 expression, CPT1 activity, diminished pyruvate and fatty acid oxidation, altered cardiolipin composition, and elevated ROS collectively depict a pre-pathological mitochondrial phenotype that likely precedes the onset of insulin resistance and chronic disease.

To our knowledge, this is the first human study to demonstrate that apparently healthy sedentary individuals exhibit a quantifiable mitochondrial transport defect centered on MPC1, detectable through non-invasive physiological biomarkers. This establishes a mechanistic framework linking molecular mitochondrial alterations to whole-body metabolic performance, redefining sedentarism not as the absence of activity but as a distinct, reversible state of early bioenergetic dysfunction.

### Mitochondrial Dysfunction and Substrate Oxidation

SED individuals showed substantial reductions across the electron transport system (Complex I −36%, Complex II −28%, ETS −34%, ATP synthase −30%), indicating a broad depression of oxidative phosphorylation capacity a reduced ability to sustain mitochondrial ATP production. This pattern aligns with classic and modern findings showing that physical inactivity decreases mitochondrial content, respiratory enzyme activity, and oxidative capacity in human skeletal muscle, as previously (Coyle et al., 1985; Fritzen et al., 2019; Houmard et al., 1992; Houston et al., 1979).

Such reductions limit the muscle’s ability to oxidize both fatty acids and carbohydrates efficiently and place greater strain on upstream metabolic pathways. This weakened oxidative machinery also sets the stage for the substrate bottlenecks described later in the discussion, particularly those involving pyruvate transport and downstream TCA flux. Together, these observations suggest that inactivity produces a coordinated decline in mitochondrial structure and function that contributes to the early development of metabolic inflexibility.

### MPC1 Reduction as a Central Mitochondrial Defect

Among all measured parameters, one of the most significant and mechanistically revealing findings was the 49% reduction in mitochondrial pyruvate carrier 1 (MPC1) expression in SED individuals, tightly correlated with a 37% decline in pyruvate oxidation. Expression of GLUT4 transporters was similar in both groups. Hence, our findings identify MPC1 as a potential primary control point for carbohydrate flux into mitochondria and establishes an “inside-out” model of glucose dysregulation, rather than impaired glucose uptake at the sarcolemma as the limitation probably arises internally at the level of mitochondrial pyruvate entry. This restriction of pyruvate transport forces glycolytic carbon to divert towards cytosolic lactate production generating a Warburg-like metabolic pattern even under aerobic conditions. Our metabolomic data reinforce this interpretation showing decreased levels of citrate and malate in SED muscle, consistent with reduced pyruvate availability to the mitochondrial matrix and impaired carbon entry into the Krebs cycle. As a consequence, cytosolic NADH accumulates, mitochondrial NAD regeneration declines and oxidative metabolism becomes progressively constrained. These events initiate a self-reinforcing cycle of redox imbalance, lactate accumulation, and diminished FATox, a pattern mirrored by the CPET data where SED participants exhibited higher lactate and lower FATox at moderate workloads.

Although cross-sectional, the multiscale convergence across MPC1, pyruvate oxidation, metabolomics, isotopologue tracing, and CPET strongly suggests that impaired mitochondrial pyruvate transport may be an early upstream defect rather than a secondary adaptive change. Mechanistically, low MPC1 expression limits the provision of acetyl-CoA to the TCA cycle, reducing electron donation to the ETS and the cell’s maximal ATP synthesis potential. This translates into a reduced capacity to oxidize carbohydrates, establishing an early subclinical state of mitochondrial insufficiency that could precede insulin resistance. Our findings therefore, provide the first experimental evidence that sedentarism is associated with markedly decreased MPC1 expression and impaired pyruvate-driven respiration, revealing a potentially reversible mitochondrial defect.

### Integrative Metabolomics and MPC1 Limitation

Metabolomic profiles separated sedentary and active individuals. Active participants displayed lower glycolytic intermediates and a balanced spectrum of acylcarnitines, a pattern consistent with coordinated mitochondrial substrate use and adequate coupling between β-oxidation, TCA cycle flux, and pyruvate oxidation. This represents a metabolically flexible phenotype in which fatty acid and carbohydrate oxidation are efficiently matched to mitochondrial demand (Muoio, 2014). In contrast, sedentary individuals accumulated pyruvate-proximal glycolytic intermediates and glyceraldehyde-3-phosphate. This pattern indicates a glycolytic bottleneck produced when pyruvate cannot enter mitochondria efficiently. Such substrate backlog is a recognized consequence of reduced MPC1 expression or activity, which limits mitochondrial pyruvate transport and forces greater reliance on cytosolic NADH reoxidation and lactate regeneration (Bricker et al., 2012; Herzig et al., 2012).

Stable isotope tracing with ¹³C-lactate confirmed this interpretation. Sedentary individuals showed markedly reduced incorporation of ¹³C into early TCA intermediates, including 40 % lower citrate labeling and 35% lower malate labeling. This pattern matches classical MPC inhibition studies where diminished pyruvate entry results in lower labeling of TCA cycle intermediates despite normal or elevated glycolytic flux (Vacanti et al., 2014).

Together, these results identify MPC1 limitation as a central mechanistic node linking glycolytic backlog, reduced mitochondrial pyruvate utilization, and the early metabolic dysfunction observed in sedentary muscle.

### Comparative Role of Fatty Acid Oxidation

In addition to pyruvate transport defect, reduced CPT1 activity and fatty acid oxidation further compound the bioenergetic limitations of the sedentary phenotype. Impaired long-chain fatty acid entry into mitochondria restricts β-oxidation and limits acetyl-CoA, reducing-equivalent (NADH, FADH) supply to the ETS. This dual restriction on carbohydrate and lipid catabolism narrows substrate flexibility and can lead to intramyocellular lipid accumulation, lipid spillover, and redox stress, which are well-established early features of metabolic inflexibility and insulin resistance as previously described by Goodpaster, Amati, Bergman and others (Amati et al., 2011; Bergman & Goodpaster, 2020; Bergman et al., 2010; Goodpaster et al., 2001)

### Dual-Substrate Bottleneck as a Hallmark of Metabolic Inflexibility

When the mitochondrial defects are considered together, a clear pattern emerges. Reduced MPC1 expression restricts pyruvate entry into the matrix, while lower CPT1 activity and impaired long-chain fatty acid transport limit β-oxidation. These two lesions converge at the level of mitochondrial carbon supply, creating a dual-substrate bottleneck that constrains both carbohydrate and lipid oxidation. With fewer carbon units entering the TCA cycle and fewer reducing equivalents delivered to the respiratory chain, oxidative phosphorylation becomes less efficient and the muscle relies increasingly on cytosolic ATP production rather than the complete oxidation of glucose through OXPHOS. This shift represents an early metabolic reprogramming toward less efficient energy generation and leaves the system more vulnerable to redox imbalance, incomplete substrate oxidation, and the early metabolic changes that can precede insulin resistance.

### Cardiolipin Remodeling and Cristae Function

Beyond its structural role, the decline in L4-cardiolipin observed in sedentary muscle may destabilize several functionally coupled components of the inner mitochondrial membrane. L4CL is required for the stability of respiratory supercomplexes and for the efficient transfer of electrons from complex I through complex III and IV. Loss or oxidation of L4CL weakens these interactions and decreases the efficiency of electron flow, which promotes ROS generation (Paradies et al., 2014; Pfeiffer et al., 2003) Higher monolysocardiolipin (MLCL) in AC suggests more active CL remodeling, supporting mitochondrial maintenance under regular physical activity (Chicco & Sparagna, 2007). Together, these mechanistic insights provide a biologically coherent explanation for how the decline in L4CL in sedentary muscle could contribute to the reduced pyruvate oxidation and electron transport capacity observed in our dataset.

### ROS, Redox Balance, and Mitohormesis

Sedentary individuals showed higher ROS production when normalized to oxygen flux, which indicates increased electron leak relative to respiratory throughput. This pattern aligns with prior observations that low mitochondrial flux and elevated membrane potential promote superoxide formation and reduce the efficiency of electron transfer (Brand 2010; Murphy 2009). Such conditions shift the redox environment toward oxidative distress rather than controlled signaling. Active participants produced higher absolute ROS, although these values scaled proportionally with their greater oxygen flux. This pattern is consistent with the concept of redox eustress, where ROS act as adaptive signaling molecules that support mitochondrial biogenesis, antioxidant gene induction, and metabolic remodeling during exercise (Gomez-Cabrera et al., 2008; Hood et al., 2011; Ristow et al., 2009). In this context, higher ROS represent a functional part of mitohormesis rather than damage. On the other hand, chronic inactivity appears to shift ROS signaling toward distress, favoring oxidative damage to mtDNA and respiratory proteins as hallmarks of early metabolic decline.

### Exercise Correlations and Translational Value

In active individuals, lactate and FATox displayed near-perfect inverse coupling (r = −0.99, p < 0.001; Fig. 10A), reflecting efficient mitochondrial function, substrate regulation, and overall metabolic flexibility. As we previously showed (San-Millan & Brooks, 2018), this pattern indicates preserved coordination between carbohydrate and lipid oxidation during incremental workloads. By contrast, sedentary individuals exhibited an early lactate inflection and a sharp decline in FATox even at mild workloads (∼100 W; Fig. 10B), denoting metabolic inflexibility and impaired mitochondrial substrate reprograming.

This phenomenon is mechanistically consistent with the inhibitory role of lactate on lipid metabolism. Elevated blood lactate levels have been shown to suppress adipose lipolysis (Liu et al., 2009), while our recent findings demonstrate that intracellular lactate also reduces CPT1 and CPT2 activity (San-Millan et al., 2022), directly limiting mitochondrial fatty acid entry and oxidation. Thus, lactate functions not only as a redox intermediate but also as an endocrine and autocrine regulator that constrains fat oxidation when chronically elevated.

CHOox and lactate increased in parallel in both groups (r ≈ 0.99, p < 0.001; Fig. 10E–F). However, in sedentary participants, the crossover point between FATox and CHOox occurred at significantly lower workloads, quantitatively demonstrating reduced mitochondrial flexibility and a premature transition toward carbohydrate dependence (Fig. 10C–D). These results indicate that the balance between lactate accumulation and FATox during graded CPET provides a direct, mechanistically interpretable index of mitochondrial substrate-oxidative capacity.

### Linking Exercise Metabolism to Mitochondrial Function

When exercise-derived variables were cross-correlated with resting mitochondrial parameters (Figs. 11–12), consistent multi-scale relationships emerged, linking cellular and whole-body bioenergetics. Exercise blood lactate at moderate intensities correlated inversely with key mitochondrial markers, including pyruvate oxidation (r = −0.57, p<0.01), palmitoylcarnitine-supported FA oxidation (r = −0.60 p<0.01), ETS capacity (r = −0.59, p<0.01), ATP-synthase-coupled flux (r = −0.62, p<0.01), and MPC1 expression (r = −0.73, p< 0.01). Exercise FATox from CPET, conversely, correlated positively with the same mitochondrial features: pyruvate oxidation (r = 0.65, p<0.01), palmitoylcarnitine flux (r = 0.71, p<0.001), ETS capacity (r = 0.63, p<0.01) ATP-synthase flux (r = 0.70, p<0.001), and MPC1 expression (r = 0.71, p < 0.001). Sarcolemmal MCT1 expression displayed weaker associations (r =0.37), indicating that intracellular oxidative capacity rather than transmembrane transport determines exercise metabolic responses in sedentary individuals.

Recent evidence indicates that a distinct pool of MCT1 resides on the inner mitochondrial membrane, functionally associated with LDH and MPC1 as part of the mitochondrial lactate oxidation complex (mLOC) (Hashimoto et al., 2006; Leija et al., 2025). The dissociation between total MCT1 protein expression and lactate oxidation rates supports the hypothesis that the limitation is not sarcolemmal uptake, but rather intramitochondrial transport and oxidation, likely involving the mLOC, which was not directly quantified here. mLOC coupling to MPC1 as part of a mitochondrial reticulum (Leija et al., 2025), provides a structural basis for the tight correlation observed between lactate flux, pyruvate oxidation, and ETS capacity in the present study.

Furthermore, lactate flux, representing the dynamic mitochondrial utilization of lactate, further reinforced this link between rest and exercise physiology (Fig. 12). Lactate flux correlated strongly and positively with pyruvate oxidation (r = 0.89, p < 0.001), MPC1 expression (r = 0.73, p < 0.05), Palmitoylcarnitine-driven FA oxidation (r = 0.95, p < 0.0001), Octanoylcarnitine oxidation (r = 0.79, p < 0.0001), ETS capacity (r = 0.84, p < 0.01) and ATP-synthase-coupled respiration (r = 0.81, p < 0.01). It correlated inversely with circulating blood lactate (r = −0.66, p = 0.05) and showed only a modest relationship with MCT1 expression (r = 0.35, p = 0.13 ns), underscoring as mentioned *vide supra* that mitochondrial uptake and oxidation, not sarcolemmal transport, constitute the main bottleneck.

Figure 11 reveals that exercise blood lactate and FATox are complementary, mechanistically grounded surrogates of mitochondrial dysfunction. Together, they integrate substrate entry, TCA flux, and electron transport into two clinically measurable outputs: ΔLactate, which reflects pyruvate transport and carbohydrate oxidation efficiency, and ΔFATox, which reflects long-chain fatty acid transport and β-oxidation capacity

### Translational Clinical Implications

The combined alterations in lactate handling and fatty acid oxidation have direct clinical value. For example, a simple observation such as blood lactate above 2.5 mmol/L together with FATox below 0.4 g per min during moderate exercise at 50 to 60 percent of VO max can serve as an early physiological signature of subclinical mitochondrial dysfunction. These values reflect impaired pyruvate oxidation and reduced long-chain fatty acid utilization, two defects that commonly appear long before overt metabolic disease develops.

The integration of these variables into a single submaximal cardiopulmonary exercise test provides a practical and scalable mitochondrial health index. This ΔLactate + ΔFATox dyad captures the two major arms of substrate oxidation. ΔLactate reflects the efficiency of glucose metabolism and the capacity of pyruvate to enter mitochondria and be oxidized through the electron transport system. ΔFATox monitors the lipid oxidative pathways linked to CPT1 function and β-oxidation. Together, they allow clinicians and researchers to identify whether the primary metabolic limitation is related to carbohydrate oxidation, lipid oxidation, or both, arising from early mitochondrial decay or dysfunction.

This approach has the advantage of being noninvasive, repeatable and sensitive to early mitochondrial impairment. It also maps directly onto therapeutic interventions, mainly through exercise. Improvements in ΔLactate + ΔFATox suggest successful restoration of substrate oxidation through an overall improvement in pyruvate and fatty acid transport and mitochondrial function. Future research should expand to include women, explore epigenetic regulation of MPC (histone lactylation, miRNA suppression), and evaluate exercise or MPC-activating interventions as reversible therapies for early mitochondrial dysfunction.

In summary, lactate and FATox responses obtained from a simple submaximal exercise test offer complementary and mechanistically grounded insight into mitochondrial efficiency. Embedding these variables into routine CPET would allow clinicians to adopt a clinically meaningful diagnostic tool for early detection of metabolic risk, personalized intervention and long term monitoring of mitochondrial and metabolic health.

### Evolutionary and Clinical Perspective

Sedentarism is not a neutral human condition but a recent biological insult. From an evolutionary standpoint, mitochondria originated as energy-producing symbionts selected under near-constant physical demand. When this demand disappears, mitochondrial structure and function undergo molecular atrophy. Our findings frame sedentarism as an early, reversible metabolic injury rather than an inevitable outcome of modern life. The convergence of molecular, metabolic and physiological evidence presented here indicates that loss of mitochondrial function is one of the first measurable steps in the trajectory toward non-communicable disease.

### Limitations and Future Directions Limitations

This study has some limitations. The sample size was modest, although it is comparable to other mechanistic human studies involving biopsies, metabolomics, and high-resolution respirometry and the data were supported by a priori power analysis. Furthermore, the large effect sizes observed across key mitochondrial variables (Cohen’s d > 1.2) which underscore the biological magnitude of these differences. However, the cohort included only men, so the findings may not generalize to women. Sedentary participants had slightly higher BMI, which may contribute to some of the metabolic differences. Finally, this was a cross-sectional comparison. Although the mechanistic logic implies that inactivity induces these mitochondrial deficits (atrophy due to disuse), longitudinal detraining and retraining studies are necessary to map the precise timeline of MPC1 restoration and metabolic recovery.

### Future directions

Future work should include women and larger, ethnically diverse cohorts. Interventional studies with structured training will help determine whether the MPC1 and CPT1 signatures are reversible and whether they improve in parallel with mitochondrial function and substrate utilization. Longitudinal designs will be important to test whether this phenotype precedes, parallels, or follows changes in insulin sensitivity. Stable isotope flux methods can be expanded to quantify tissue-specific lactate oxidation and its regulation by MPC1 and redox state. Finally, external validation of physiological markers from submaximal CPET could allow development of a clinical tool for early mitochondrial health assessment.

## Conclusion

Healthy sedentary adults display a coordinated impairment in mitochondrial metabolism that includes lower MPC1 abundance, reduced pyruvate and lipid oxidation, altered cardiolipin structure, and diminished lactate oxidation. These molecular and organelle-level features are reflected in characteristic shifts in fat oxidation and lactate accumulation during CPET, which provides a practical non-invasive window into underlying mitochondrial function. Together, these findings support the presence of an early and potentially reversible mitochondrial phenotype associated with inactivity. The integrated profile described here offers a set of physiological and molecular markers for future longitudinal and interventional studies and highlights MPC1 and related mitochondrial components as promising targets for strategies aimed at restoring oxidative capacity.

## Acknowledges

We thank the University of Colorado Anschutz Medical Campus for facilities, technical staff, and participants. Funded by internal ISM grants.

## Author Contributions

ISM: conceptualization, supervision, exercise testing, data analysis, manuscript writing, funding acquisition. JLM: exercise testing, data analysis. GCS: cardiolipin analysis. AD, DS, TN: metabolomics and fluxomics. JH: muscle biopsies. All authors reviewed the manuscript.

## Data Availability

Raw data (metabolomics, respirometry, exercise) will be available upon publication. Contact the corresponding author for interim access.

